# Accuracy, speed and error tolerance of short DNA sequence aligners

**DOI:** 10.1101/053686

**Authors:** Mark Ziemann

**Affiliations:** Epigenetics in Human Health and Disease Laboratory, Baker IDI Heart and Diabetes Institute, The Alfred Medical Research and Education Precinct, Melbourne, Victoria 3004

**Keywords:** High throughput sequencing, Short read aligners, DNA alignment

## Abstract

Aligning short DNA sequence reads to the genome is an early step in the processing of many types of genomics data, and impacts on the fidelity of downstream results. In this work, the accuracy, speed and tolerance to errors are evaluated in read of varied length for six commonly used mapping tools; BWA aln, BWA mem, Bowtie2, Soap2, Subread and STAR. The accuracy evaluation using Illumina-like simulated reads showed that accuracy varies by read length, but overall BWA aln was most accurate, followed by BWA mem and Bowtie2. BWA mem was most accurate with Ion Torrent-like read sets. STAR was at least 5 fold faster than Bowtie2 or BWA mem. BWA mem tolerated the highest density of mismatches and indels compared to other mappers. These data provide important accuracy and speed benchmarks for commonly used mapping software.

## Introduction

The development and adoption of high throughput sequencing (HTS) in the past 15 years had brought about a new era in genomics. Illumina sequencing in particular has been a major contributor to public DNA sequence databanks and continues to be the platform of choice for large scale genomics projects **[1,2](Abecasis et al, 2012; Peplow 2016)**. Aligning short reads to the genome is an early data processing step and impacts the quality of downstream findings.

There are a range of sequencing applications suited to short or longer reads. For example, relatively short reads of 50 to 100 nt are commonly used to map epigenetic modifications with ChIP-seq. However, exome and whole genome and metagenome sequencing commonly utilises paired-end format with read lengths of 100 nt or longer. Reported maximum read lengths! are 250 nt for HiSeq2500 Rapid and 300 nt for MiSeq. As paired end mode is available in all these, length of merged pair sequences of up to 600 nt can be obtained **[3,4](Illumina 2015)**.

Other HTS platforms are also available including semiconductor sequencing on Ion Torrent systems **[5](Rothberg et al, 2011)**. The popular Ion PGM instrument generates reads with a range of lengths from 35 to 400 nt **[6](Thermo Fisher 2013)**. Ion Torrent sequence data has a relatively high rate of indel errors compared to Illumina systems, as a result of inability to accurately distinguish homopolymers **[7](Loman et al, 2012)**.

Genomic alignment has its challenges, particularly the presence of sequence polymorphisms, genomic repeats, and the large size of some genomes such as human. Accuracy in terms of precision (specificity) and recall (sensitivity) are important in genomics applications to limit false negative and false positive findings. Aligners must therefore be aware of genomic repeats, robust to sequence polymorphisms and efficient enough so that the huge volumes of DNA sequence data can be processed in a reasonable computational time.

Several informative evaluations have previously been undertaken to determine the most accurate alignment softwares **[8–14](Holtgrewe et al, 2011; Hatem et al, 2013; Shang et al, 2014; Caboche et al, 2014; Otto et al, 2014; Highnam et al, 2015; Smolka et al, 2015)**, but relatively few of these evaluate read lengths >100 nt or included newer aligners such as BWA mem and STAR. Few evaluations specifically determine the robustness of mappers to varied rates of mismatch and indel errors at a range of read lengths. There are also very few published analyses of the performance of mappers with Ion Torrent data.

In this work, I quantify the accuracy of six free and open source aligners with realistic simulated Illumina and Ion Torrent sequences at different read lengths in Arabidopsis (~135 Mbp genome) and human (~3.1 Gbp). The speed and robustness to sequence errors including mismatches, insertion and deletions is also evaluated. These findings will be relevant for many investigators looking to optimise short read mapping procedures.

## Methods

### Alignment accuracy of Illumina and Ion Torrent-like reads

Illumina like reads were generated using the simulator ART v2.3.7 **[15](Huang et al, 2012)**at a range of read lengths (50, 100, 200 nt) using the default built-in error profile. The 480 nt read sets were prepared by first generating paired-end 250 nt reads with ART, followed by merging with PEAR v0.9.8**[16](Zhang et al, 2014)**. Arabidopsis and human reference genomes were downloaded from Ensembl (http://asia.ensembl.org/info/data/ftp/index.html Arabidopsis_thaliana.TAIR10.30.dna.genome.fa & Homo_sapiens.GRCh38.dna.primary_assembly.fa). Aligner versions and command lines used to perform read alignments are given in Table 1. Ion Torrent reads were simulated using dwgsim v0.1.11 **[24](Homer 2011)** at read lengths of 50, 100, 200 and 480 nt and otherwise default parameters. TMAP was only evaluated for Ion Torrent-like reads.

**Table 1.**
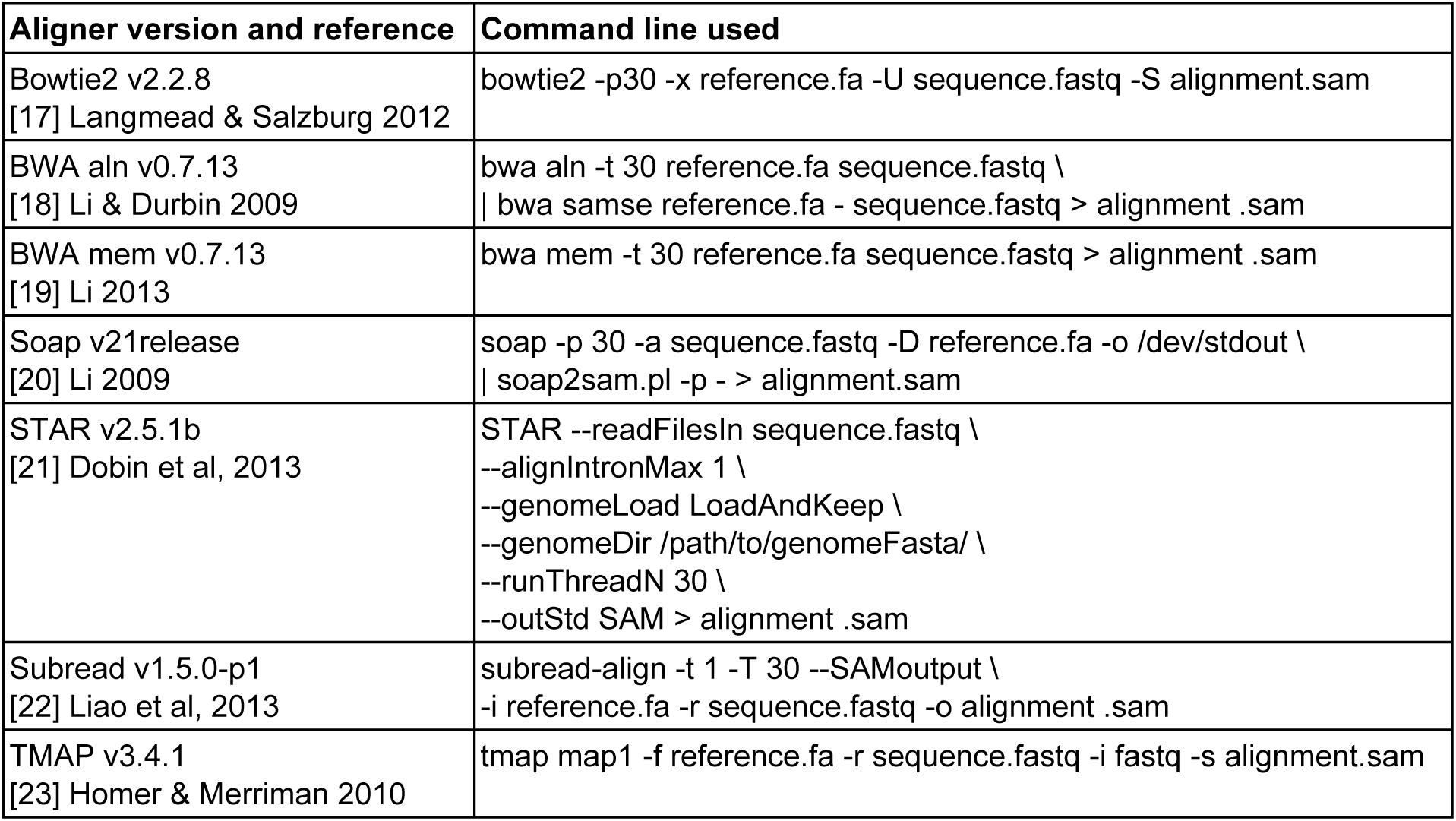
**Aligner versions and parameters used in the evaluations**.

A read is classified as correctly mapped if the start and end of the read are placed within 100 bp of the ground truth. A read is classified unmapped if there is no reported alignment or the map quality value is below the specified threshold. A read is classified as incorrectly mapped if the reported alignment is above the specified threshold with start and end coordinates >100 bp from that anticipated from ground truth. With Illumina like sequences, ground truth positions were obtained from the SAM file generated by ART. Reads generated by dwgsim or the custom script (described below), contained ground truth positions in the sequence header information. The accuracy of mappers with each test was quantified using the F-measure that considers both precision and accuracy. Throughout this paper, we use a beta value of 0.25 that weights precision higher than recall, as described previously **[25](Ziemann et al, 2016)**.

### Aligner speed evaluation

The speed of aligners was assessed by determining the time taken to process read sets of 1 million (M) and 5 M reads. Using this approach it is possible to calculate the time taken to (i) load the index into memory and (ii) calculate the speed at which reads are processed. Simulator derived reads with Illumina-like error profiles were generated with ART **[15](Huang et al, 2012)**as above. Uncompressed fastq sequences were mapped with parameters allowing up to 30 parallel threads, sending SAM-formatted alignment data to the null device (/dev/null). A custom BASH script was used to quantify computational time and resource utilisation on a 32 core 128 GB RAM Dell PowerEdge server running Ubuntu 12.04.

### Alignment accuracy of error rich reads

A custom script was used to generate perfectly matching reads in fasta format, followed by error incorporation using msbar (EMBOSS v6.4.0.0) **[26](Rice et al, 2000)**. These reads were mapped and precision/recall determined as above. All scripts to generate read sets, perform alignment and read evaluation are uploaded to SourceForge (https://sourceforge.net/projects/ziemann-dnaaligner-evaluation/).

## Results

### Accuracy of mappers with Illumina like reads of various length

In order to ascertain which aligners are most accurate for read lengths currently generated by Illumina systems, read sets with lengths 50, 100, 200 and 480 bp were generated from Arabidopsis and human templates, mapped to the respective genomes and the number of correct and incorrectly mapped reads was quantified. Overall accuracy was evaluated using the F-measure. As the F-measure is dependent on the applied mapping quality (mapQ) threshold, we calculated F-measure at a range of mapQ values for 50 bp reads (**Figures 1a**,**b**) and demonstrate that a mapQ threshold of 10 was appropriate for these six mappers. This mapQ threshold is used throughout the paper. The precision, recall and F-measure values for Arabidopsis and human read sets for reads of length 50, 100, 200 and 480 nt are shown in **Figures 1c**,**d**. In both Arabidopsis and human tests, Subread had the lowest precision, but highest recall. Precision, recall and F-measures were similar between BWA aln, BWA mem and Bowtie2. Soap2 was as precise as other mappers but had the lowest recall, especially with longer read lengths. Despite being designed specifically to map RNA sequences, STAR performed relatively well, with above average recall. Overall, recall was increased markedly in 100 nt reads as compared to 50 nt. For most mappers, recall for 200 nt reads was only slightly higher than 100 nt reads. Surprisingly, most mappers showed lower recall for 480 nt reads as compared to 200 nt, with BWA mem the only exception. The overall average F0.25 scores (including Arabidopsis and human) were determined, with Subread scoring lowest (0.969) followed by Soap2 (0.989), STAR (0.992), then Bowtie2 (0.994) and BWA mem and BWA aln tied in top place (0.995). These data quantify the accuracy of commonly used mappers with Illumina-like reads at a range of read lengths in the context of a small and large genome.

**Figure 1.**
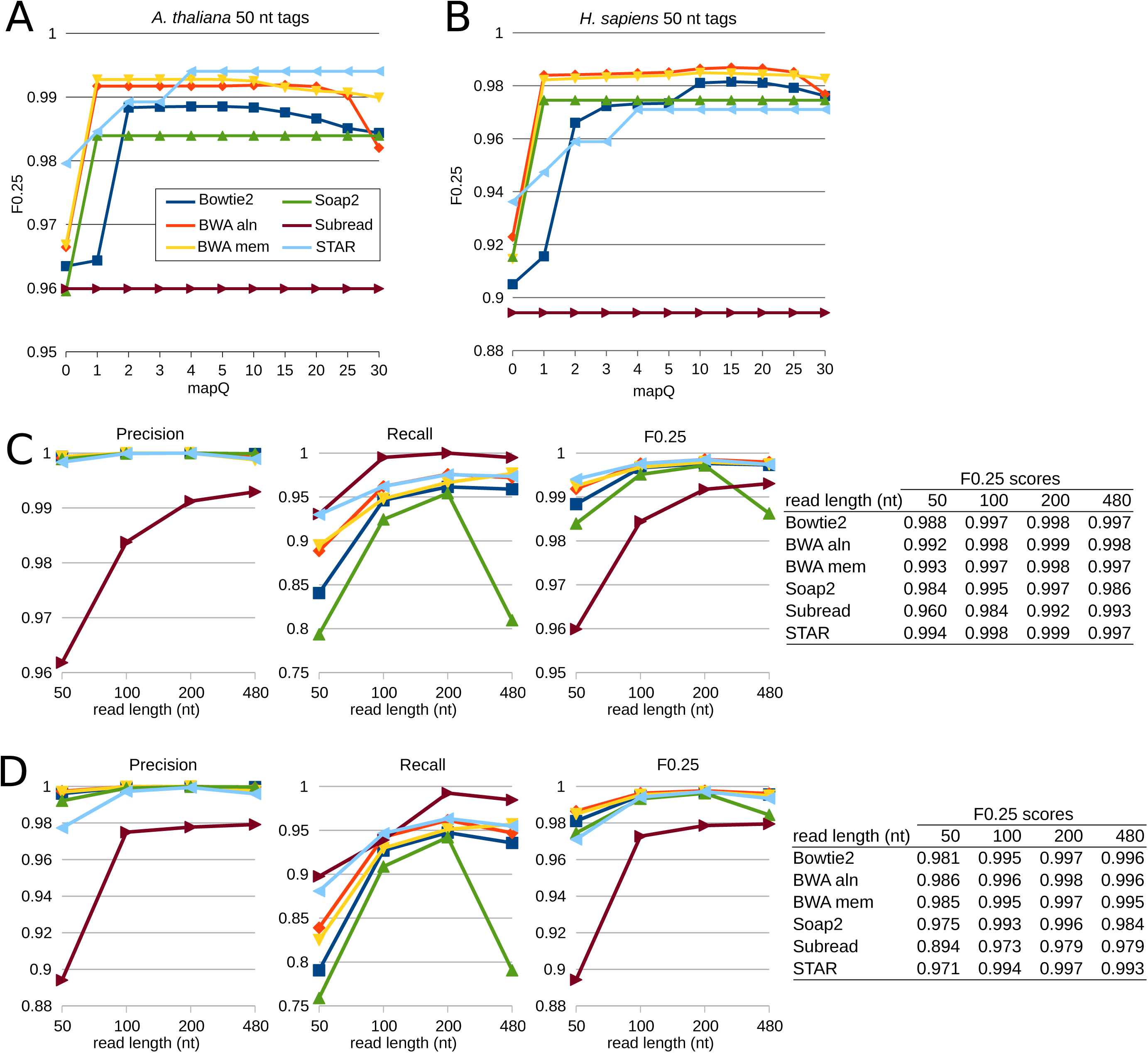
Aligner accuracy using Illumina-like reads at a range of lengths. Simulated reads were mapped to the respective genome. The F0.25 measure for 50 nt tags was calculated at a range of mapQ values for *A. thaliana* (A) and *H. sapiens* (B). MapQ threshold of 10 was used for subsequent calculation of precision, recall and F0.25 at a range of read lengths for *A. thaliana* (C) and *H. sapiens* (D).

### Accuracy of mappers with Ion Torrent like reads of various length

In order to investigate which mappers were best for processing Ion Torrent sequence reads, we performed read simulation using the DWGSIM tool **[24](Homer 2011)**to generate Ion Torrent like read sets with lengths of 50, 100, 200 and 480 nt. These were mapped using the same aligners as above with the additional inclusion of TMAP1, that is recommended by the instrument manufacturer. The precision, recall and F-measures were determined for Arabidopsis and human read sets (**Figure 2a**,**b**). Overall Subread had the lowest precision. Soap2 scored the lowest recall. BWA aln scored relatively poorly as compared to other aligners, and recall declined with increasing read length. The overall average F0.25 scores (including Arabidopsis and human) were determined, with Soap2 scoring lowest (0.504) followed by BWA aln (0.682), Subread (0.896), TMAP1 (0.971), STAR (0.976), Bowtie2 (0.984) and BWA mem (0.992) in top place. These data quantify the accuracy of commonly used mappers with Ion Torrent-like reads at a range of read lengths in the context of a small and large genome.

**Figure 2.**
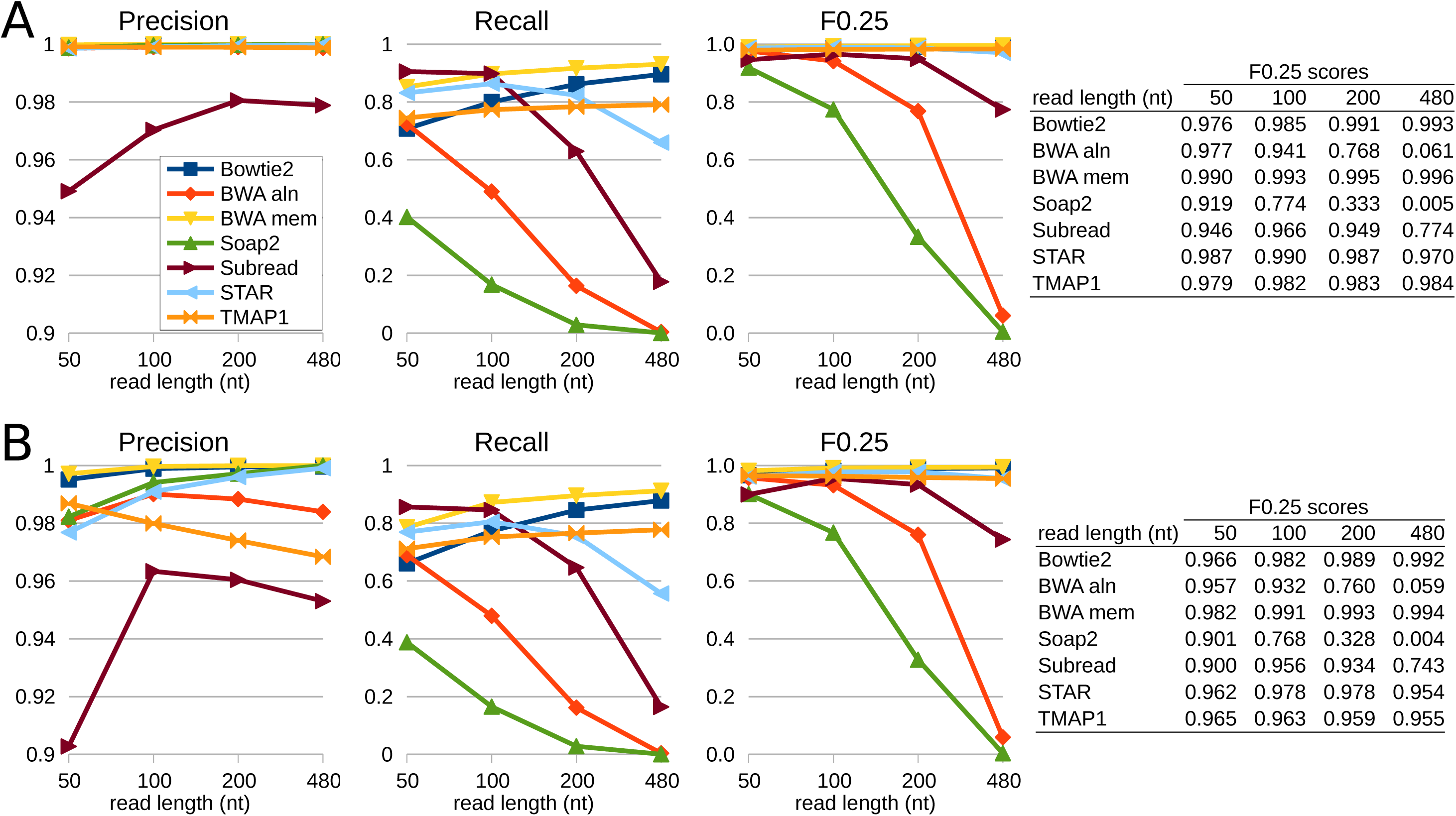
Aligner accuracy using Ion Torrent-like reads at a range of lengths. Simulated reads were mapped to the respective genome. MapQ threshold of 10 was used for subsequent calculation of precision, recall and F0.25 at a range of read lengths for *A. thaliana* (A) and *H. sapiens* (B).

### Aligner speed

In light of the increasing volume of data being produced from high throughput sequencers, alignment speed may be a factor in the selection of a mapper. The throughput at which reads were processed on a standard server allowing up to 30 threads was determined using Illumina-like read sets as above in Arabidopsis and human (**Figure 3**). STAR aligner was the fastest in all tests. In human 50 nt reads, STAR was 30.4x faster than Bowtie2 and 186x faster than BWA aln. Overall, Soap2 was the slowest in both Arabidopsis and human tests. Despite the smaller genome size, Arabidopsis alignments were, in some cases, not much faster than the human counterpart. While BWA aln ran 2.6x faster in Arabidopsis than human, STAR was actually 15% faster in human as compared to Arabidopsis. In general, increasing read length lead to slower processing time, with Bowtie2 showing the greatest slowdown (46x) as compared to BWA mem (5.5x) when comparing processing speed at 50 and 480 nt read lengths in human. These results demonstrate that STAR might be a useful alternative to researchers for high throughput DNA sequencing; future versions of STAR could incorporate built in parameter settings for DNA alignments.

**Figure 3.**
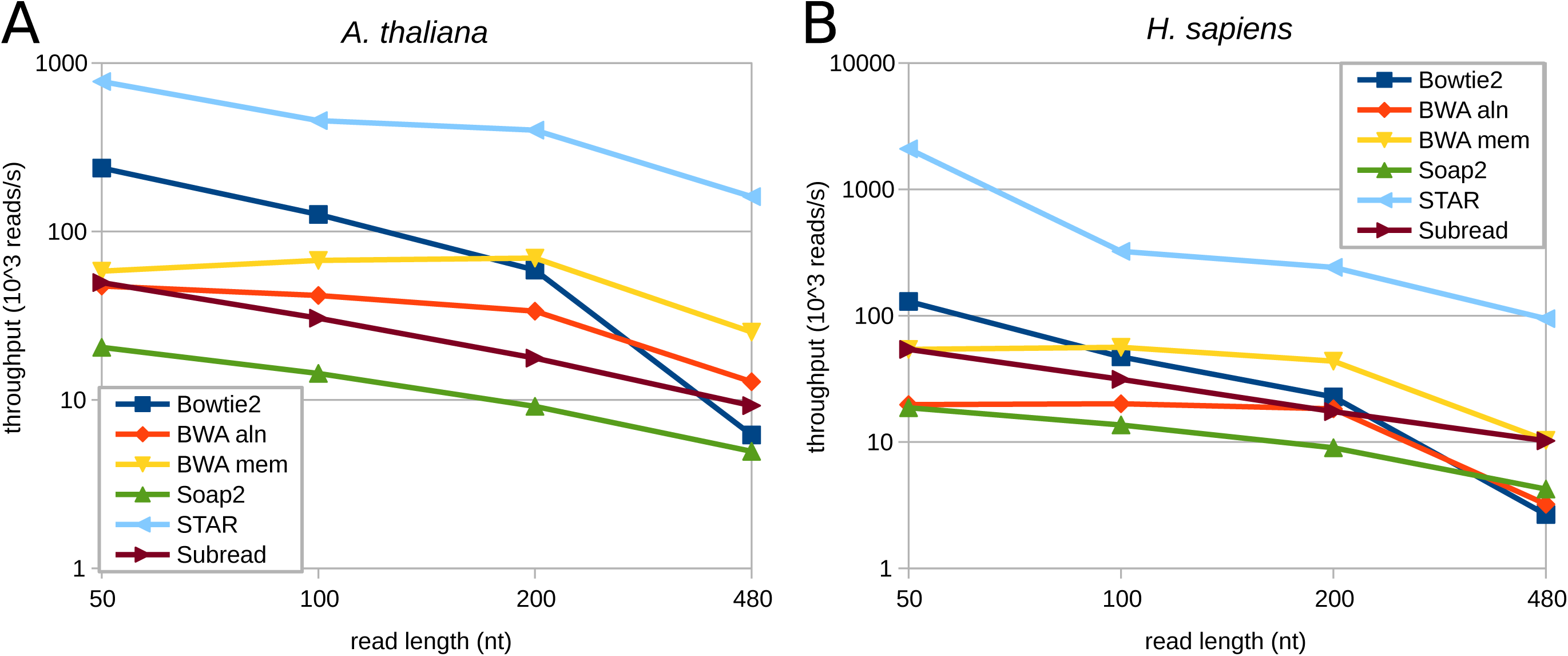
Aligner speed using Illumina-like reads at a range of lengths. (A) *A. thaliana*. (B) *H. sapiens*. Values are the median of three replicates.

### Robustness with regards to mismatches

Mismatches are common in all sequencing data and may be the result of sequencing errors or genetic variation. Accurate mapping of mismatch containing reads enables better resolution of genetic differences and reduces the number of unmapped and unused reads. To quantify the robustness with regards to single nucleotide mismatches (SNM), perfectly matching read sets were first generated, followed by incorporation of up to 16% SNMs in reads with length 50 to 500 nt. After mapping, the F-measure was determined and plotted as a function of SNM frequency (**Figure 4**). BWA mem was the most robust to SNM in reads ≥100 nt, recording F measures >0.9 even with 16% SNMs. Bowtie2 and STAR were the next best, followed by BWA aln then Subread and lastly Soap2. The tolerance of BWA aln and Soap2 to SNM diminished with read length >100 nt, whereas BWA mem was more tolerant to SNM in longer reads. With 50 nt reads, STAR was the most accurate with SNM rich reads. These results demonstrate that BWA mem accuracy is superior to other mappers with mismatch rich reads of 100 nt or longer.

**Figure 4.**
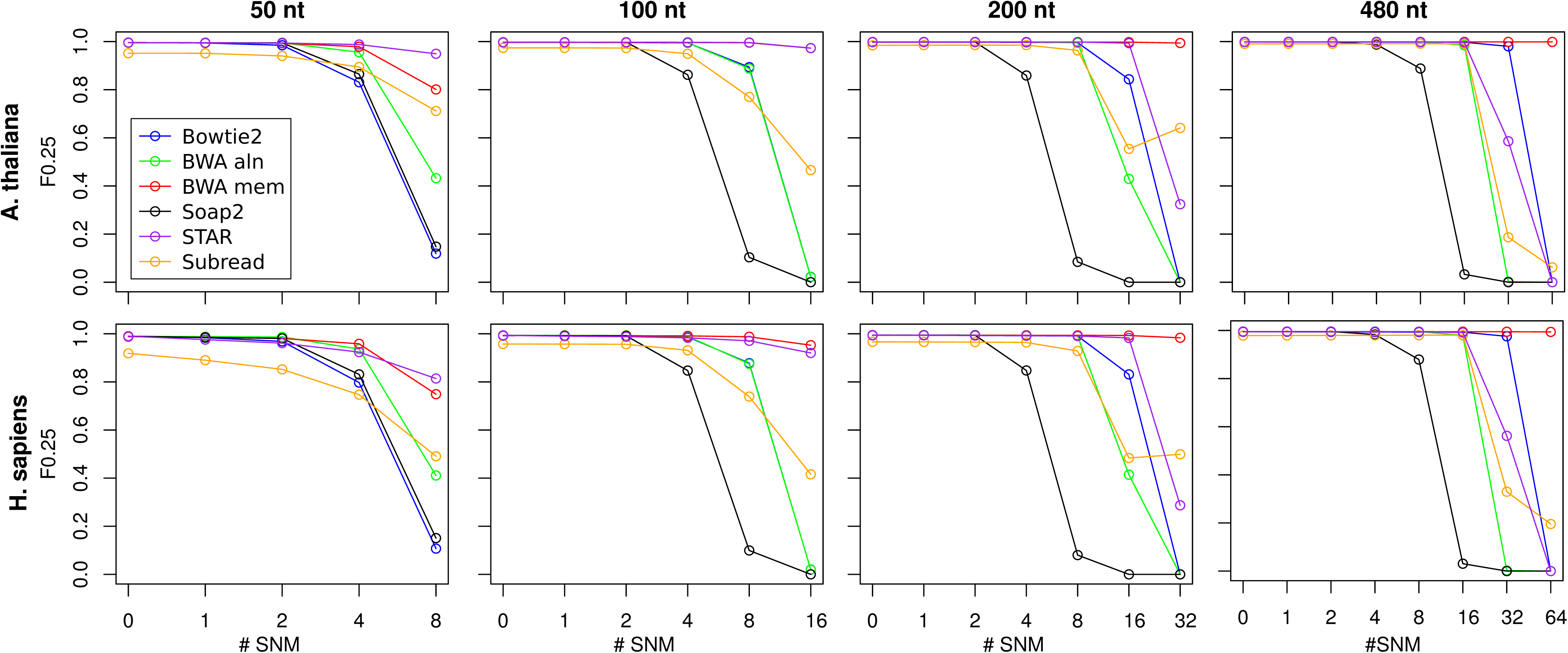
Performance of aligners with reads containing single nucleotide mismatches. The F0.25 measure is used as a quantitative measure of precision and recall.

### Robustness with regards to indels

Indel errors can be relatively common in sequencing data, depending on instrument type. For instance the Ion Torrent and Roche 454 machines yield indel rates much higher than Illumina systems due to differences in sequencing chemistry **[7](Loman et al, 2012)**. Indels also occur randomly by mutation, accumulating at 1/8th the rate of mismatches according to studies of the human Y chromosome **[27](Wei et al, 2013)**. To assess mapper tolerance to indels, insertions or deletions were incorporated into reads at a frequency up to 16%. The mapping results for indel containing human reads are shown in **Figure 5**. Results for Arabidopsis were virtually identical (not shown). Overall, the effect of indel incorporation was more deleterious to accuracy as compared to SNM. The effect of insertions and deletions was quite similar, and this was consistent in Arabidopsis and human tests. Soap2 was least tolerant to indels, followed by BWA aln. Bowtie2, Subread and STAR were moderately tolerant to indels, while BWA mem was clearly the most tolerant to indels.

**Figure 5.**
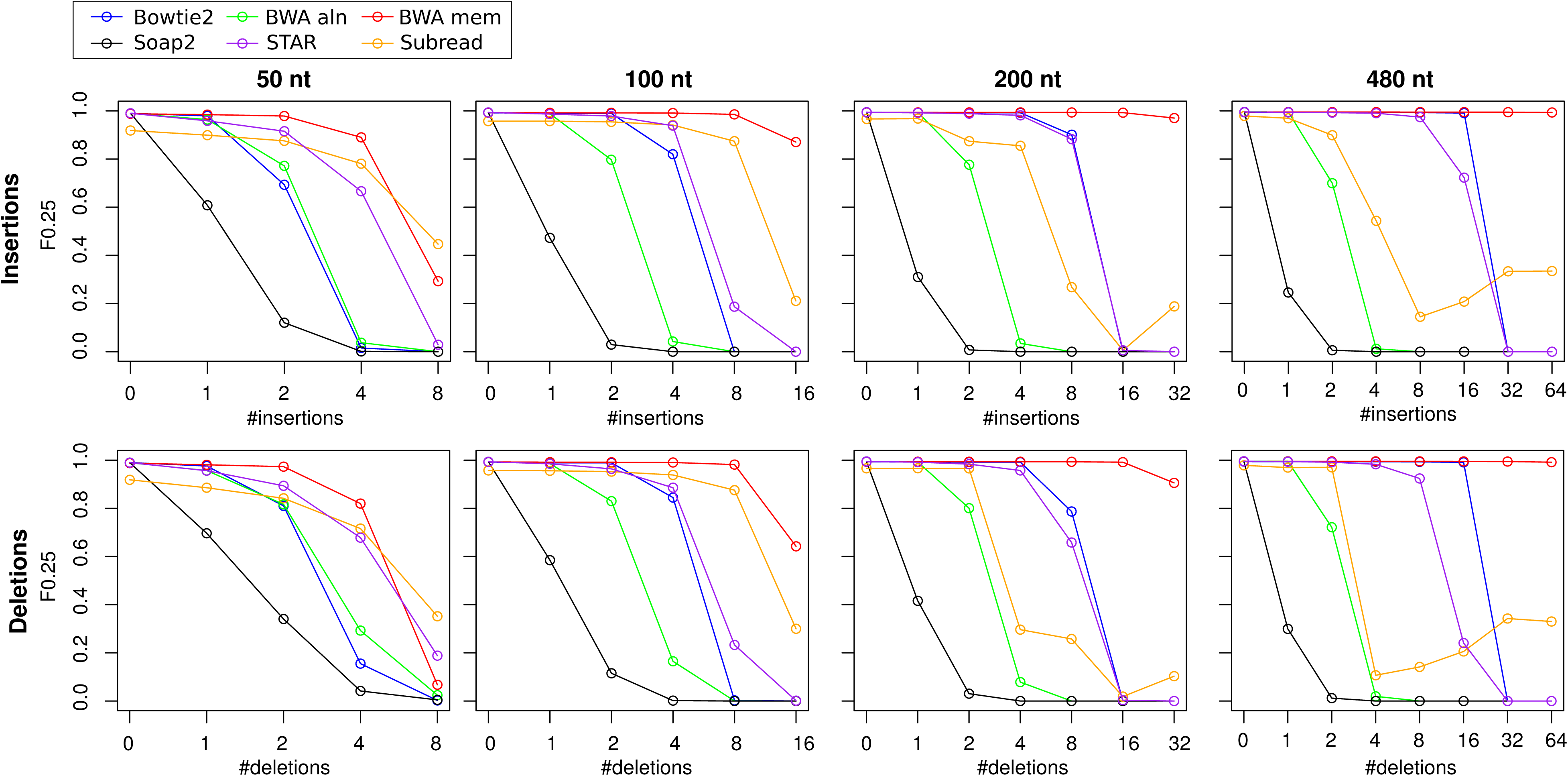
Performance of aligners with human derived reads containing single base insertions and deletions.

## Discussion

There are a multitude of short read mappers available **[28](Fonseca et al, 2012)** and the choice of mapper can be daunting for new investigators. In this work, the accuracy of six aligners with Illumina-like reads of varying length was determined. The results indicate that using a human template and standard Illumina error profiles lacking genomic variants, BWA aln is the most accurate in most situations, followed closely by BWA mem and Bowtie2. In Arabidopsis, STAR is as accurate as BWA aln, BWA mem and Bowtie2. STAR aligner is also the fastest mapper evaluated, making it a potentially useful tool for DNA mapping given the appropriate parameter settings.

Evaluation of Ion Torrent like read sets reveal BWA mem is most accurate aligner evaluated at all read lengths in Arabidopsis and human, even superior to TMAP1, the aligner recommended by the instrument manufacturer. These results indicate that BWA mem will perform well for Roche 454 read sets, as Ion Torrent and Roche 454 data have similar error profiles **[7](Loman et al, 2012)**.

The mappers evaluated showed a broad range of tolerances to mismatches and indels. Overall Soap2 and BWA aln are most sensitive to error incorporation and as such, tightly clustered SNM and indel variants could be missed by these tools. In contrast, BWA mem is highly robust to SNMs and indels in reads of 100 nt or longer.

These findings show that BWA mem is highly accurate and tolerant to errors including mismatches and indels. These features make BWA mem suited to mapping of whole genome bisulfite sequencing reads. Indeed the BWA mem algorithm has already been harnessed to perform this task and performs well in simulations **[29](Pederson et al, 2014)**.

Taken together, this work outlines the strengths and weaknesses of several commonly used open source short read mapping tools for DNA alignment.

